# Combination CDC-like kinase inhibition (CLK)/Dual-specificity tyrosine-regulated kinase (DYRK) and taxane therapy in *CTNNB1*-mutated endometrial cancer

**DOI:** 10.1101/2023.04.04.535570

**Authors:** Bradley R Corr, Marisa R Moroney, Elizabeth Woodruff, Zachary L Watson, Kimberly R. Jordan, Thomas Danhorn, Courtney Bailey, Rebecca J Wolsky, Benjamin G Bitler

## Abstract

SM08502 (cirtuvivint) is a novel pan CDC-like kinase (CLK) and Dual specificity tyrosine kinase (DYRK) inhibitor that targets mRNA splicing and is optimized for Wnt pathway inhibition. Previous evaluation of single agent CLK/DYRK inhibition (SM04690) demonstrated inhibition of tumor progression and β-catenin/TCF transcriptional activity in *CTNNB1*-mutant endometrial cancer (EC). *In-vitro* analysis of SM08502 similarly decreases Wnt transcriptional activity and cellular proliferation while increasing cellular apoptosis. SM08502 is an active single-agent therapy with IC50’s in the nanomolar range for all EC cell lines evaluated. Combination of SM08502 with paclitaxel has synergistic effect *in vitro*, as demonstrated by Combination Index <1, and inhibits tumor progression in four endometrial cancer models (HEC265, Ishikawa, Ishikawa-S33Y, and SNGM). In our *in vivo* mouse models, Ishikawa demonstrated significantly lower tumor volumes of combination vs SM08502 alone (Repeated Measures one-way ANOVA, p = 0.04), but not vs paclitaxel alone. HEC265, SNGM, and Ishikawa-S33Y tumors all had significantly lower tumor volumes with combination SM08502 and paclitaxel compared to single-agent paclitaxel (Repeated Measures one-way ANOVA, p = 0.01, 0.004, and 0.0008, respectively) or single-agent SM08502 (Repeated Measures one-way ANOVA, p = 0.002, 0.005, and 0.01, respectively) alone. Mechanistically, treatment with SM08502 increases alternative splicing (AS) events compared to treatment with paclitaxel. AS regulation is an important post-transcriptional mechanism associated with the oncogenic process in many cancers, including EC. Results from these studies have led to a Phase I evaluation of this combination in recurrent EC.

## INTRODUCTION

Endometrial cancer (EC) is the most common gynecologic malignancy in the developed world, and both the incidence and mortality of EC are on the rise worldwide. Over the last decade in the United States, EC incidence rose from 42,190 new cases in 2009 to 61,880 new cases in 2019, while estimated annual deaths rose from 7,780 to 12,160^1, 2^. EC is one of the only cancers to have a raising mortality rate, and with this increasing trajectory of both incidence and mortality, EC will likely soon be responsible for more annual deaths in the United States than ovarian cancer^1–5^.

Recurrent metastatic endometrial cancer often portends a poor prognosis^6^. With advancements in molecular tumor analysis guiding patient-centered precision medicine, there have been recent treatment successes for the population of patients with recurrent EC. The Cancer Genome Atlas (TCGA) initially identified subgroups of EC with distinct genetic profiles and statistically different outcomes^7^. The prognostic significance of these subgroups has subsequently been independently reproduced and verified, ultimately demonstrating the clinical implications of a molecular based risk-stratification system^8–11^. Molecular based treatment strategies are now commonly used in practice. Examples include the use of PD-L1 blockade (pembrolizumab or dostarlimab) for recurrent mismatch repair deficient/microsatellite instable (MMRd/MSI-H) recurrent endometrial cancers^12, 13^; the use of combination pembrolizumab and lenvatinib for mismatch repair proficient (MMRp) recurrent endometrial tumors^14^, and the use of trastuzumab in carcinomas that overexpress Her2/Neu^15^.

*CTNNB1* mutations have been identified as an important clinical marker in EC^7–10, 16^. *CTNNB1* is a gene that codes for the β-catenin protein, which is involved in the Wnt/β-catenin pathway and is associated with multiple cancers^7, 16–21^. In normal cells in the absence of a Wnt ligand, a destruction complex binds to and phosphorylates the N-terminus of β-catenin, ultimately resulting in the degradation of β-catenin. In contrast, when Wnt ligand is present, Wnt proteins initiate a signaling cascade that prevents β-catenin phosphorylation and degradation. β-catenin then accumulates in the cytosol, translocates to the nucleus and interacts with T cell factor/lymphoid enhancing factor (TCF/LEF) transcription factors and co-activators (e.g. cAMP response element binding protein, CREBBP) to initiate transcription^16, 18, 20–22^. *CTNNB1* mutations associated with EC are mainly located in exon 3 of the *CTNNB1* gene. Exon 3 encodes the N-terminus of β-catenin; these mutations therefore disrupt phosphorylation of β-catenin by the destruction complex, resulting in aberrant accumulation of β-catenin and subsequent transcriptional hyperactivation^8, 16, 23^.

Wnt/β-catenin signaling is involved in the regulation of the normal endometrium, and aberrant Wnt/β-catenin activity, like that with *CTNNB1* mutations, has been associated with the development of endometrial hyperplasia and malignancy ^16, 24–26^ ^25, 27^. Multiple studies of ECs with otherwise low mutational burden report that tumors that contain *CTNNB1* exon 3 mutations are more likely to recur^7, 8^. Moreover, *CTNNB1* mutations have been demonstrated to be associated with a significantly increased rate of disease recurrence specifically in a low-risk population of early stage, low grade ECs^28^. In larger studies evaluating ECs of all grades, ECs with *CTNNB1* exon 3 mutations have been found to have decreased recurrence-free and overall survival^8, 23, 28, 29^. Combined, this data indicates the CTNNB1 mutations are potential oncogenic drivers.

We demonstrated that downstream inhibition of the Wnt/β-catenin pathway can decrease cell viability, inhibit TCF/β-catenin transcriptional activity, and diminish growth in endometrial *in vitro* and *in vivo* models^30^. Our previously published work evaluated SM04690 (which is a small molecule that indirectly inhibits TCF/β-catenin transcriptional activity by inhibiting intranuclear CDC-like kinase 2 (CLK2) and thus disrupting alternative splicing^31^. Based on the mechanism of action of SM04690, we confirmed that transcriptional activity would be diminished with its use. The compound SM04690 is not bioavailable, but a next generation bioavailable drug labelled SM08502 is in clinical development. SM08502 is a small molecule inhibitor of Cdc2-like kinases (CLK1-4) and the dual-specificity tyrosine phosphorylation regulated kinases (DYRK1-4) that modulates alternative mRNA splicing and is optimized for Wnt pathway inhibition^32^.

Single agent paclitaxel is an effective, and well tolerated, treatment modality for recurrent endometrial cancer^33–35^. The combination strategy of SM08502 and taxanes originated from previous work evaluating Wnt inhibitors and taxanes. Fischer *et al.* demonstrated synergistic effect in combining the Wnt inhibitors vantictumab and ipafricept with paclitaxel by potentiating mitotic cell death^36^. This combination has been evaluated in multiple non-gynecologic phase IB studies demonstrating tolerability and promising efficacy^37, 38^.

In this report, we define the efficacy of SM08502 in both *in vitro* and *in vivo* models of *CTNNB1*-mutated endometrial cancer. Furthermore, the combination of SM08502 and paclitaxel were assessed in a similar panel of endometrial cancer models. We also determined the SM08502-dependent transcriptome and using bioinformatic tools we measured alternative-splicing changes. We demonstrate that the SM08502 and paclitaxel combination strategy is synergistic in preclinical models and can be an effective strategy for patient evaluation.

## MATERIALS AND METHODS

### RNA Extraction

FFPE tissue blocks were acquired (COMIRB#20-1067) and a board-certified pathologist confirmed tumor tissue. This protocol is deemed exempt, as it is using previously collected data, and the information is not recorded in a manner that is identifiable. FFPE tissue blocks were sectioned into 10 micron tissue-containing paraffin scrolls. RNA was extracted using the High Pure FFPET RNA Isolation kit (Roche). RNA quantity and quality was assessed using a RNA Screentape on a TapeStation 4150 (Agilent). RNA concentration was determined by comparison to the RNA ladder and the percentage of RNA fragments greater than 200 bp was calculated (average 70.6%, all samples > 55%).

### NanoString PanCancer Immuno-Oncology (IO) 360

150 ng of RNA was combined with hybridization buffer and the Reporter CodeSet for the PanCancer IO 360 Panel (Nanostring) and incubated for 20 h at 65°C. The hybridized reaction was analyzed on an nCounter SPRINT Profiler (Nanostring). nSolver calculated normalization factors for each sample using raw gene counts and 14 housekeeping genes. Differential gene expression was calculated from the normalized gene counts and false discovery rates with a Benjamini Hochberg multi-comparison test were determined. The average count for the negative control probes was used for thresholding “positive” genes. After thresholding (<20 counts), a total of 666 genes were subsequently used for downstream analysis. The heatmap was generated using Clustergrammer^39^. Raw gene counts are normalized using the logCPM method, filtered by selecting the genes with most variable expression, and transformed using the Z-score method.

nSolver advanced analysis tool was used to generate a pathway score for 25 different pathways (e.g., Hypoxia). The pathways scores were grouped and compared based on *CTNNB1* mutant vs *CTNNB1* wild type status. Genetic signature analysis was performed by Nanostring as previously described^40–43^. The log2-transformed gene expression values are multiplied by pre-defined weighted coefficient^40^ and the sum of these values within each gene set is defined as the signature score.

### Cell Culture

Ishikawa (RRID: CVCL_2529), HEC265 (RRID: CVCL_2928), and SNGM (RRID: CVCL_1707) are human EC cell lines. Ishikawa is *CTNNB1*-wildtype, while HEC265 and SNGM are *CTNNB1*-mutant (*CTNNB1* Exon 3 base substitutions D32V and S37P, respectively). Ishikawa-S33Y is a *CTNNB1* mutated cell line created by retroviral transduction of the S33Y-mutant CTNNB1 gene (pBabe-CTNNB1-S33Y) into the Ishikawa cell line^30^. All four cell lines were authenticated using small tandem repeat analysis (The University of Arizona Genetics Core), and the *CTNNB1* status of each cell line was confirmed through Sanger Sequencing. The HEC1B and HEC265 cells were cultured in Minimum Essential Medium (MEM) medium supplemented with 1% penicillin-streptomycin, and 15% fetal bovine serum. Ishikawa cells were cultured in MEM medium supplemented with 1% penicillin-streptomycin, 1% non-essential amino acids, 1% glutamine, and 5% fetal bovine serum. SNGM cells were cultured in 1:1 DMEM and Ham’s F12 supplemented with 20% FBS and 1% penicillin-streptomycin. All cells were maintained in 5% CO_2_ at 37°C and were routinely tested for mycoplasma with MycoLookOut (Sigma-Aldrich).

### Reagents

SM08502 was obtained from Biosplice Therapeutics (San Diego, CA 92121). Paclitaxel was obtained from Millipore Sigma (Cat. No. Y0000698).

### Dose Curves and Crystal Violet Staining

The effect of SM08502 on cell viability was evaluated in the EC cell lines via crystal violet staining. Each cell line was plated in a 96-well plate with 10,000 cells per well (n = 4). The cells were incubated for 48 hours following treatment. Crystal violet staining was then performed to measure cell viability with cell survival normalized to control (0.1% DMSO) being the measure for cell viability. Briefly, after treatment cells were fixed (10% methanol, 10% acetic acid) and stained with 0.4% crystal violet. Crystal violet was dissolved in fixative, and absorbance was measured at 570 nm on a SpectrMax plate reader.

### Combination Index

Cells were added to 96 well plates at a concentration of 10,000 cells per well and were placed in an incubator overnight to allow cells to reach confluency. Cells were then treated with SM08502 or paclitaxel alone or in combination at varying concentrations. Plates were placed back in an incubator for 48 hours after which time cells were stained with 0.4% crystal violet. Then, the cells were fixed (10% methanol, 10% acetic acid) and stained with 0.4% crystal violet. Crystal violet was dissolved in fixative and absorbance was measured at 570 nm. Combination index analysis was performed using CompuSyn for Drug Combinations software^44^.

### Apoptosis Assay

Cells (n = 3, each cell line) were plated in a six-well culture dish and allowed to attach for 24 hours. The cells were then treated with vehicle or SM08502. After 48 hours of treatment, the cells were stained with Alexa Fluor 488 annexin V kit (Cat A13201; Invitrogen) and propidium iodide according to the manufacturer’s protocol. The cells were then analyzed using a Gallios Flow Cytometer at the University of Colorado Cancer Center Flow Cytometry Facility. FlowJo (v10) was used to analyze data.

### TCF Transcriptional Reporter

TCF transcriptional activity was evaluated using a Luciferase Assay System (Cat. E1501; Promega) and TOP-FLASH, FOP-FLASH plasmids. TOP-FLASH was a gift from Randall Moon (Addgene plasmid # 12456; http://n2t.net/addgene:12456; RRID:Addgene_12456). FOPFlash (TOPFlash mutant) was a gift from Randall Moon (Addgene plasmid # 12457; http://n2t.net/addgene:12457; RRID:Addgene_12457). Using FuGENE6 reagent (Cat. E2692; Promega), populations were transfected with TOP-FLASH or FOP-FLASH plasmid. Cells were incubated for 24 hours, then moved to a 96-well plate and treated with serial doses of SM08502 (n = 4, each cell line). Following treatment, the cells were incubated for another 48 hours. Cells were then lysed and analyzed using the Luciferase Assay system with luminescence measured by a Promega GloMax.

To normalize this assay for transfection efficiency and cell count, FOP-FLASH luciferase activity and crystal violet staining were performed on each cell line. For FOP-FLASH transfected cells, luminescence was also quantified as a measure of transfection efficiency. For crystal violet staining, the cells were seeded, incubated and treated in the same way as the cells in the Luciferase Assay System. Then, 48 hours after treatment, the cells were fixed (10% methanol, 10% acetic acid) and stained with 0.4% crystal violet. Crystal violet was dissolved in fixative and absorbance was measured at 570 nm.

### Anti-BrdU FITC Cell Staining and Flow Cytometry

Cells (n = 3, each cell line) were plated in 6-well culture dishes and incubated in the appropriate growth media for 24 hours at 37°C and 5% CO_2_. The cells were then treated with either vehicle or SM08502 (600, 1200, 1800, or 2400 nmol/L) for 48 h. 5′-bromo-2′-deoxyuridine (BrdU) (Cat. #550891; BD Biosciences; RRID:AB_2868906) was then added directly to the well culture media (final concentration 10 µM), and the cells were incubated at 37°C for 60 minutes. After BrdU incorporation, cells were washed twice with phosphate-buffered saline (PBS) and treated with 0.25% trypsin/0.1% EDTA for 7 minuts at 37°C followed by two washes with 1% bovine serum albumin (BSA)/PBS and resuspension in cold PBS. Finally, cells were slowly added to ice cold 70% ethanol and incubated at −20°C for 30 minutes.

Fixed cells were incubated for 30 minutes in 2 N HCl with 0.5% Triton X-100 (vol/vol) followed by resuspension in 0.1 mol/L Na_2_B_4_O_7_·10 H_2_O, pH 8.5. Cells were then suspended in 0.5% Tween 20 (vol/vol) plus 1% BSA/PBS and incubated for 30 minutes at room temperature with anti-BrdU FITC (BD Biosciences, Cat. #347583) at a concentration of 0.5 µg/10^6^ cells. Lastly, cells were washed once in 0.5% Tween 20 (vol/vol) plus 1% BSA/PBS and resuspended in PBS. The anti-BrdU FITC cells were then analyzed using a Gallios Flow Cytometer at the University of Colorado Cancer Center Flow Cytometry Facility. Laser excitation was set at 488 nm. FlowJo (v10) was used to analyze data.

### RNA sequencing

Cells (n = 2, each cell line) were plated in 6-well culture dishes and incubated with the appropriate growth media for 24 hours at 37°C and 5% CO_2_. They were then treated with either vehicle control (0.1% DMSO) or their respective IC_50_ concentrations of paclitaxel (Ishikawa: 10.9 nM, Ishikawa-S33Y: 6.1 nM, HEC265: 5.6 nM, SNGM: 11.7 nM), SM08502 (Ishikawa: 63.4 nM, Ishikawa-S33Y: 79 nM, HEC265: 134.4 nM, SNGM: 350.7 nM) alone or in combination. After 24 hours treatment, RNA was extracted using RNeasy columns with on-column DNAse treatment (Qiagen, Cat. # 74004). Ribosomal RNA depletion was performed using QIAseq FastSelect -rRNA kit (Qiagen, Cat. #335376). 350 ng input RNA was used for Ishikawa cells; 500 ng input RNA was used for SNGM, HEC265, and S33Y cells. Step 1 of fragmentation was performed for 7 minutes at 94°C, followed by FastSelect hybridization. First-strand synthesis, second-strand synthesis, A-tailing, adapter ligation, and library amplification were performed using the KAPA mRNA HyperPrep kit (Roche, Cat. #08105952001). We used 3.33 μL of 1.5 μM KAPA single index adapters for Illumina (Roche, Cat #0800577001) for adapter ligation. We performed 11 amplification cycles. KAPA Pure magnetic beads (Roche, Cat. #08105901001) were used for all cleanup steps. Library quality was confirmed by TapeStation by the University of Colorado Cancer Center Pathology Shared Resource. Sequencing was performed on a NovaSeq6000 instrument by the University of Colorado Genomics and Microarray Core.

For differential gene expression FASTQ files were aligned to human genome (Ensembl annotation release 87). HISAT2^45^ was used for alignment against GRCh37 version of the human genome. Samples were normalized using TPM (Transcripts per Million) measurement and gene expression using the GRCh37 gene annotation was calculated using home-made scripts. The analysis was performed utilizing BioJupies^46^. RNA-sequencing data has been deposited to NCBI GEO: GSE215975.

### Splicing Analysis

For alternative-splicing analyses, BBDuk (version 38.90, https://sourceforge.net/projects/bbmap/) was used to remove Illumina TruSeq adapters and remove sequence reads with a remaining length of less than 50 nt. Reads were aligned to the GRCh38 assembly of the human genome with the STAR mapper (version 2.7.9a)^47^ in end-to-end alignment mode, using splice junction information from Ensembl release 104 (https://may2021.archive.ensembl.org/Homo_sapiens/). Alternative splicing events were detected and compared between treatments within each cell line using rMATS (version 4.1.2)^48^ with transcript annotations from Ensembl release 104. Events detected using both junction counts and exon body counts (JCEC) with a false discovery rate (FDR) of 0.05 or less were counted for each of the five categories (skipped exon, retained intron, mutually exclusive exons, as well as alternative 5’ and 3’ splice sites). The authors acknowledge a potential bias in the analysis technique in that rMATS only evaluate annotated retained introns events. As previous retained intron events would be captured in the annotations, it is possible that those captured are more prone to retention. Analysis was conducted by the University of Colorado Anschutz Medical Campus Cancer Center Biostatistics and Bioinformatics Shared Resource (BBSR) core facility (RRID:SCR_021983).

### Mouse Models

All mouse work was approved under a University of Colorado Institutional Animal Care and Use Committee (IACUC) protocol. Athymic nude mice (Charles River Labs, Strain 553) were subcutaneously injected with 5 × 10^6^ HEC265, Ishikawa, or Ishikawa-S33Y cells, or 1 × 10^7^ HEC265 or SNGM cells on the right flank. Tumor progression was measured using calipers every 2 days, and tumor volume as well as % change in tumor volume ((day x volume – day 0 volume)/day 0 volume) x 100) were recorded. Measurements were used to determine tumor volume based on the formula a^2^ x b/_2_, with “a” being the smaller diameter and “b” being the larger diameter. Once a tumor in each flank population grew to over 100 mm^3^, treatment was initiated in all mice. For all experiments, mice were treated with paclitaxel alone (10 mg/kg weekly, i.p.), SM08502 alone (12.5 mg/kg, daily, PO), paclitaxel + SM08502, or vehicle. Body weight was measured twice per week during treatment as a surrogate for toxicity. Body weight loss of >10% was considered indicative of severe reaction to treatment. The mice were sacrificed according to IACUC protocol, and the tumors were surgically resected and weighed. Tumor burden was calculated based on the weight of resected tumors. Blood samples were collected at time of death via cardiac punction for complete blood count (CBC) and blood chemistry analysis (AST, ALT, and ALP), blood volume permitting. Blood samples were analyzed at the University of Colorado Comparative Pathology Shared Resource.

### Ki67 Immunohistochemistry methods

Tumor tissue from mouse model specimens were fixed in 10% buffered formalin and stored in 70% ethanol. The tumor tissue was then paraffin-embedded and sectioned for hematoxylin and eosin staining and immunohistochemistry (IHC). IHC staining was performed for Ki67 (Cat. #RM-9106-S, 1:200; Thermo Fisher Scientific; RRID:AB_2341197). This tissue preparation, sectioning and staining was performed by the Histopathology Core of The University of Colorado Cancer Center as previously described^49^. All slides were deidentified and imaged at 20x magnification on a Olympus CKX41 microscope. The images were imported and analyzed in QuPath to calculate the percentage of Ki67+ cells.

### Statistical Analysis

All statistical analyses were performed in Prism GraphPad (v9). Dose curve calculations were performed using nonlinear regression (variable slope), and the IC_50_ of each drug for each cell line was determined. Standard statistical analyses with one-way ANOVA and Repeated Measures one-way ANOVA were performed with Tukey’s post hoc analysis, where indicated. A p<0.05 or adjusted p<0.05 were considered significant.

### Data Availability

Cell line data are presented in Supplemental table 1. All other data are available upon request from the corresponding author

## RESULTS

### Genomic variation is present in human patient CTNNB1 mutant vs wild type recurrent stage I endometrial cancer tumors

Patients with recurrent stage I, grade 1 endometrioid adenocarcinoma of the endometrium were evaluated for *CTNNB1* mutational status and demonstrated to have increased risk of recurrence based on our previous case control study^28^. We identified 5 CTNNB1 mutant vs 4 wild type tumors from this patient subset for transcriptomic analysis. FFPE tumor tissue was identified and utilized for RNA isolation and transcriptomic analyses. We employed the NanoString platform to examine the expression of 770 highly annotated genes including housekeeping genes in the PanCancer IO 360 panel, which targets genes expressed by tumor cells and immune cells in the tumor microenvironment^43^. Figure 1 A&B demonstrate the heat map and volcanic plot illustrating a clear separation of gene expression of the two subgroups. Figure 1 C&D illustrate the variation in known genomic pathways indicating that the three pathways with the most significant variation are Wnt signaling, angiogenesis, and the PI3K-AKT pathways.

**Figure 1:**
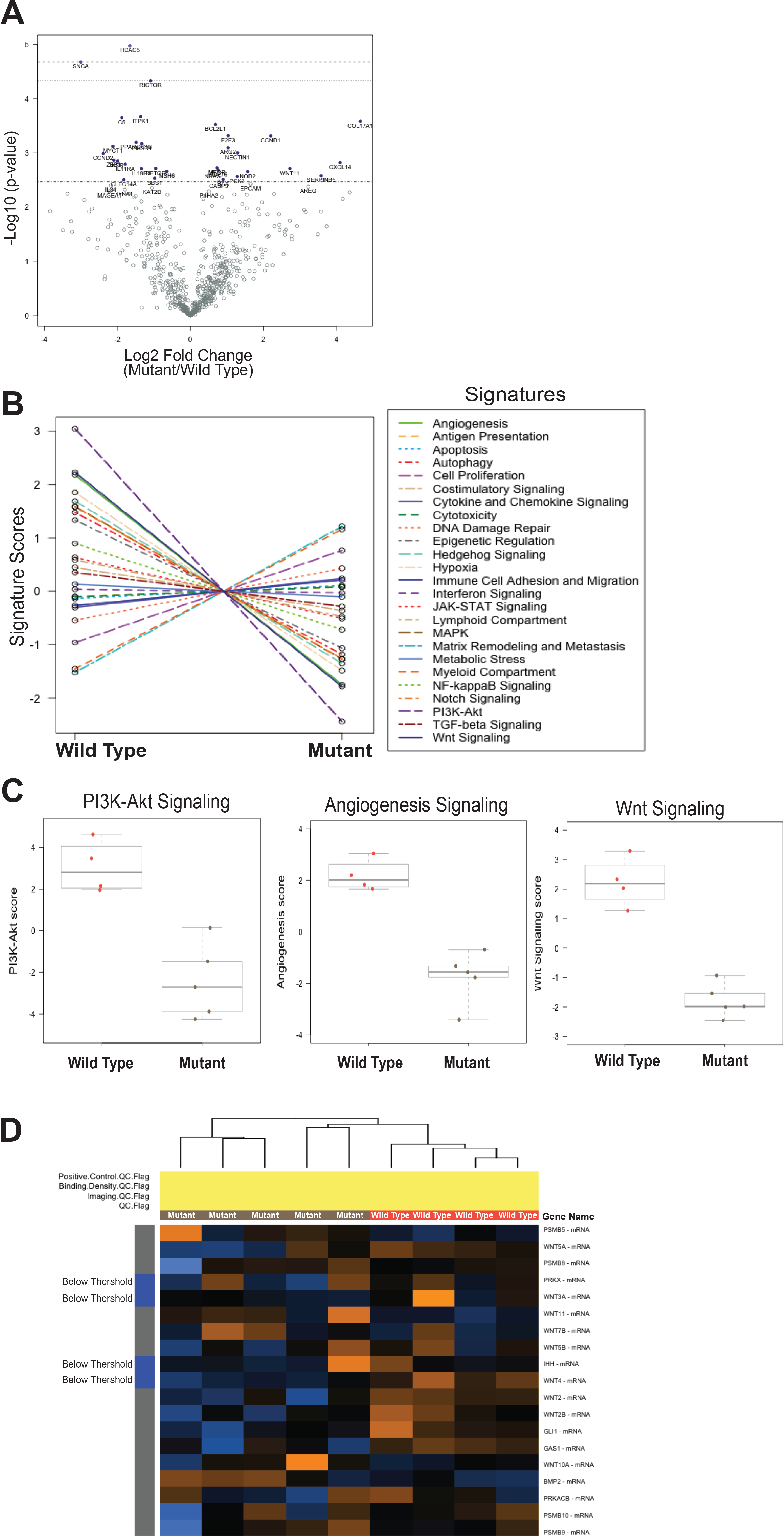
NanoString transcriptomic analysis of *CTNNB1* mutant vs wild type recurrent stage I grade 1 endometrioid adenocarcinomas of the endometrium. **A)** Volcano plot of differentially regulated genes. P-value calculated with paired t-test with Benjamin-Hochberg multi-comparison correction (Adj. p-value). **B)** Pathway scores condense each sample’s gene expression profile into a set of predetermined pathway scores. Pathway scores are fit using the first principal component of each gene set’s data. They are oriented such that increasing score corresponds to mostly increasing expression (specifically, each pathway score has positive weights for at least half its genes). The pathway scores are plotted to show how they vary across *CTNNB1* mutant vs wild type conditions. Lines show each pathway’s average score across values of mutant. **C)** Plots of determined (Wnt signaling, angiogenesis, and PI3K-Akt) pathway scores for *CTNNB1* mutant vs wildtype. **D)** Heatmap of the 19 differentially expressed Wnt pathway genes, generated with Clustergrammer.

### SM08502 is a first in class pan CDC/DYRK inhibitor that decreases TCF transcription, increases apoptosis, and downregulates cellular proliferation

Single agent therapy against four distinct EC cell lines shows uniformly decreased cellular survival with IC50’s in the nanomolar range, regardless of *CTNNB1* mutational status (Figure 2A). TCF transcriptional activity was measured using TOP-FLASH/FOP-FLASH luciferase-based reporter system. Compared to control, single-agent SM08502 (800 nM) significantly decreased TCF transcriptional activity in all four cell lines regardless of CTNNB1 status (HEC265: -93.9 %TCF activity, p < 0.0001; SNGM: -45.9%, p = 0.03; Ishikawa-S33Y: -93.3%, p < 0.0001; Ishikawa: -93.1%, p < 0.0001) (Figure 2B). We examined the effect of SM08502 on apoptosis and proliferation via Annexin V/PI and BrdU incorporation, respectively. Compared to control, 1200 nM SM08502 significantly induces apoptosis in Ishikawa (7.3% vs. 16.63% AnnexinV/PI+, p < 0.0001), HEC265 (12.47% vs. 39.9% AnnexinV/PI+, p < 0.0001), and SNGM (19.93% vs. 67.3%, p < 0.0001), but not Ishikawa-S33Y (Figure 2C). A correlation with *CTNNB1* status is not readily apparent. With respect to apoptosis, a dose response is apparent in the Ishikawa and SNGM cell lines starting at a dose of 600 nM. In contrast a static significant increase in apoptosis is seen in the HEC265 cell line starting at a dose of 1200 nM. With respect to proliferation, SM08502 significantly inhibited BrdU incorporation in all three *CTNNB1*-mutant cell lines (HEC265, SNGM, and Ishikawa-S33Y), while having no significant impact on proliferation of *CTNNB1*-wild type Ishikawa cell line (Figure 2D).

**Figure 2:**
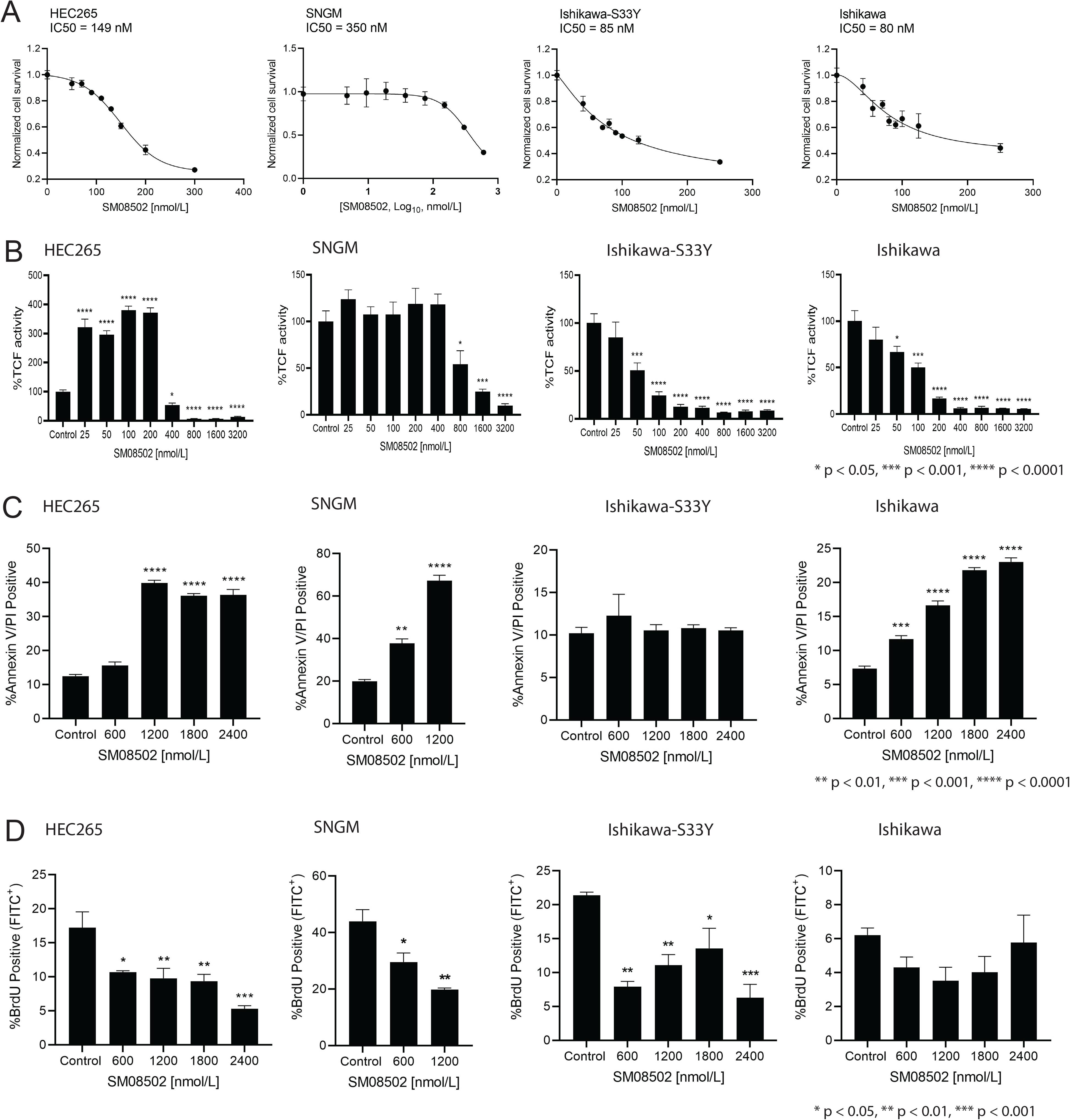
Dose response and β-Catenin/TCF transcriptional activity in CTNNB1-wildtype and CTNNB1-mutant cell lines following treatment with SM08502. **(A)** Dose response curves for CTNNB1-wildtype (Ishikawa) and CTNNB1-mutant (HEC265, SNGM, Ishikawa-S33Y) cell lines performed by measuring %cell confluency with crystal violet stain following treatment with serial doses of SM08502 (nmol/L). **(B)** TCF transcriptional activity measured via TOP-FLASH luciferase-based reporter system. Cell lines were transfected with either TOP-or FOP-FLASH and treated with increasing doses of SM05802 for 48 h. Luminescence was normalized to cell number and FOP-FLASH signal (transfection control). **(C)** Apoptosis measured by % cells Annexin V/PI positive following serial dosing of SM08502 **(D)** Cells treated with SM08502 for 48h and incubated with BrdU for 1h. Experiments conducted at least two independent times in triplicate. Statistical test, one-way ANOVA, Tukey multicomparison. *p < 0.05, **p < 0.01, ***p < 0.001, ****p < 0.0001. Error bars, SEM. ANOVA, analysis of variance; TCF, T cell factor

### Combination of SM08502 and paclitaxel is synergistic in vitro

In vitro combination of fixed ratio combinations of SM08502 and paclitaxel demonstrate synergy in three of the four cell lines: HEC265, Ishikawa-S33Y, Ishikawa (Figure 3A & 3B). The HEC 265 cell line demonstrates the greatest synergy with combination index (CI) scores reaching less than 1 at only 25% effect levels. Both the Ishikawa and the Ishikawa-S33Y cell lines demonstrate CI less than 1 at the higher effect levels, both roughly around the 50% effect. The SNGM cell line is the only cell line to not show synergy of the combination, however an additive effect is seen uniformly at all effect levels with a CI range between 1.15 and 1.31.

**Figure 3:**
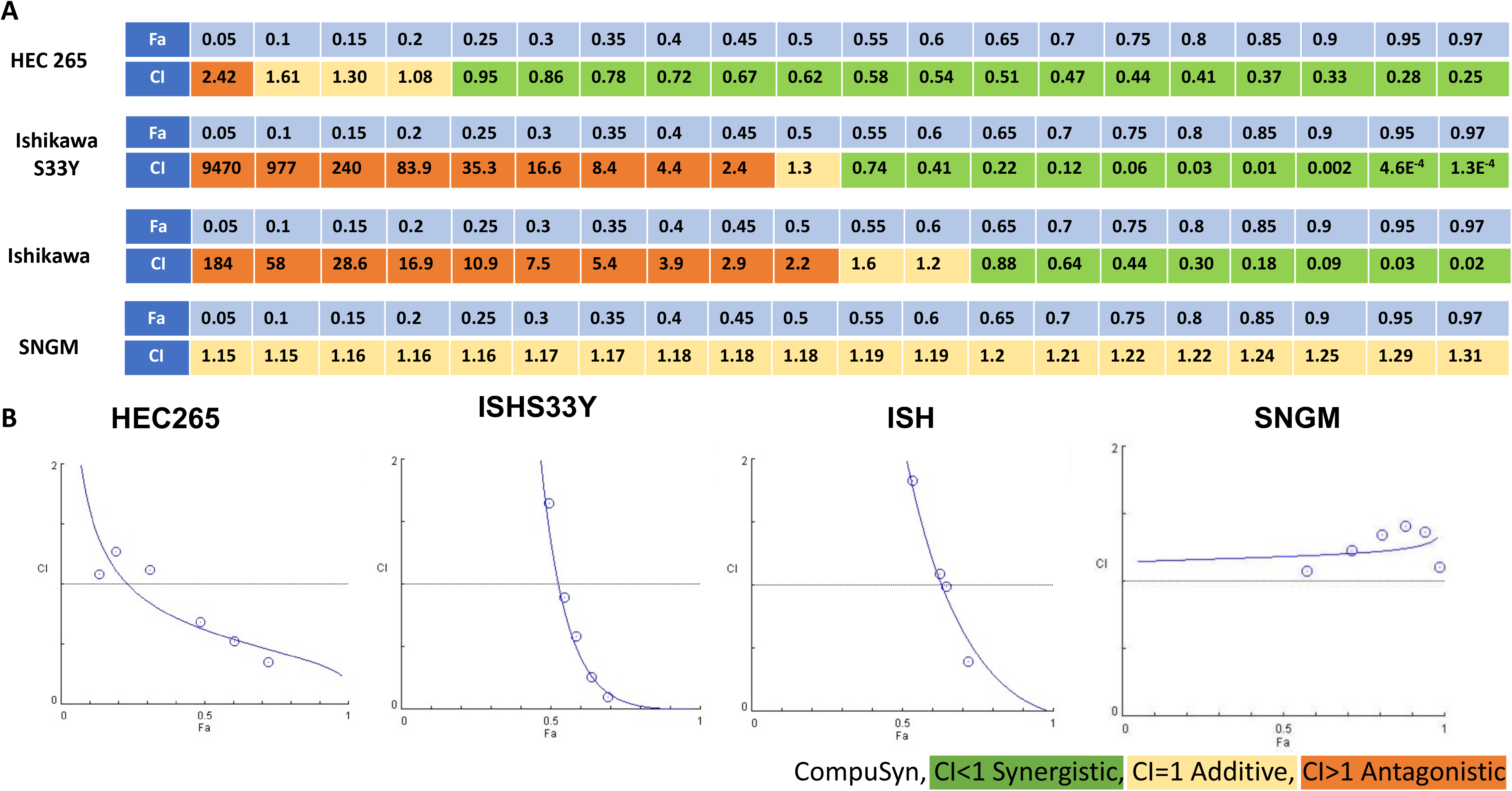
Combination index (CI) evaluation for synergy of combination SM08502 and paclitaxel. Cells were seeded into 96-well plates at a concentration of 10,000 cells per well and treated with serial concentrations of SM08502, paclitaxel, or combined SM08502/paclitaxel based on previously acquired IC50 values. Control wells were treated with 0.1%DMSO only. After 48 hours, cells were stained with crystal violet. Combination index values were determined based on crystal violet absorbance (570 nm). Compusyn (ComboSyn, inc.) programing was utilized to calculate **(A)** CI values and **(B)** graphs depicted. Calculated effect levels (Fa) are reported per Compusyn calculations.

### In vivo treatment of combination SM08502 and paclitaxel has reduced tumor growth compared to paclitaxel or SM08502 alone

Uniformly across all four cell line xenograft models (HEC265, Ishikawa, Ishikawa-S33Y, & SNGM), combination paclitaxel + SM08502 exhibited a significant reduction in tumor volume compared to control (p = 0.0023, p = 0.0055, p = 0.0012, p = 0.0031, respectively). Tumor volume was also reduced with combination treatment when compared to paclitaxel alone in HEC265, Ishikawa-S33Y, & SNGM (p = 0.0113, p = 0.0008, p = 0.0041, respectively) but not Ishikawa (p = 0.0808). Tumor volume was reduced in combination treatment compared to SM08502 alone in HEC265, Ishikawa, Ishikawa-S33Y, & SNGM (p = 0.002, p= 0.0389, p = 0.0144, p = 0.0053, respectively) (Figure 4).

**Figure 4:**
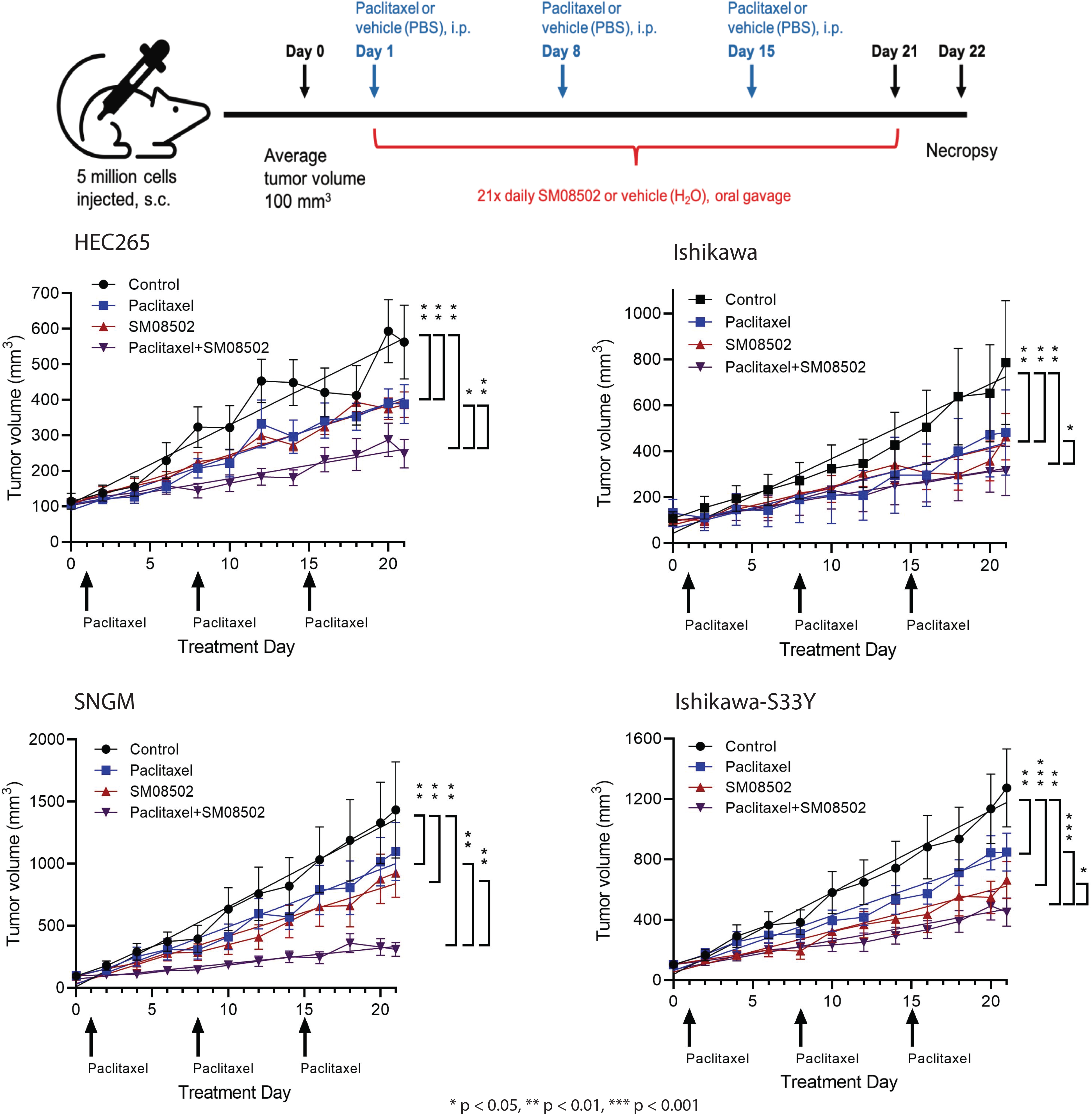
In-vivo models for treatment of combination SM08502 & paclitaxel, SM08502, paclitaxel and control. Nude mice were subcutaneously injected with either 5 (Ishikawa and Ishikawa-S33Y) or 10 (HEC265 and SNGM) million tumor xenograft cells in their right flank. Once tumors reached an average volume of 100 mm^3^, mice were treated daily via oral gavage with SM08502 (12.5 mg/kg) or vehicle (DI water) and weekly with paclitaxel (10 mg/kg) or vehicle (PBS) via IP injection for 21 days. Over the course of the 21-day study, tumors were measured with calipers every two days, and tumor volume was calculated based on the formula a^2^ × b/2 with “a” being the smaller diameter and “b” being the larger diameter. On the 22^nd^ day of the study, mice were euthanized, and tissue samples were collected. Linear regression. Error bars, SEM. Statistical test 2-way ANOVA (mixed model effect) * <0.05, ** p <0.01, *** p < 0.001

SM08502 single agent therapy was also uniformly effective in tumor reduction when compared to control (p = 0.0092, p = 0.0071, p = 0.0008, p = 0.003, respectively). Furthermore, single agent SM08502 performed with relatively equivalent efficacy as paclitaxel alone.

IHC analysis of post treatment tumors demonstrates statistically significant reduction in %Ki67+ cells with combination therapy compared to SM08502 alone in the Ishikawa-S33Y (P=0.0164) and SNGM (p=0.0157). Combination therapy also decreased %Ki67+ cells compared to paclitaxel (p=0.0041) and control (p=0.0008) in SNGM. No variation in %Ki7+ cells was seen in the HEC265 or Ishikawa cell lines (Figure 5 A-D).

**Figure 5:**
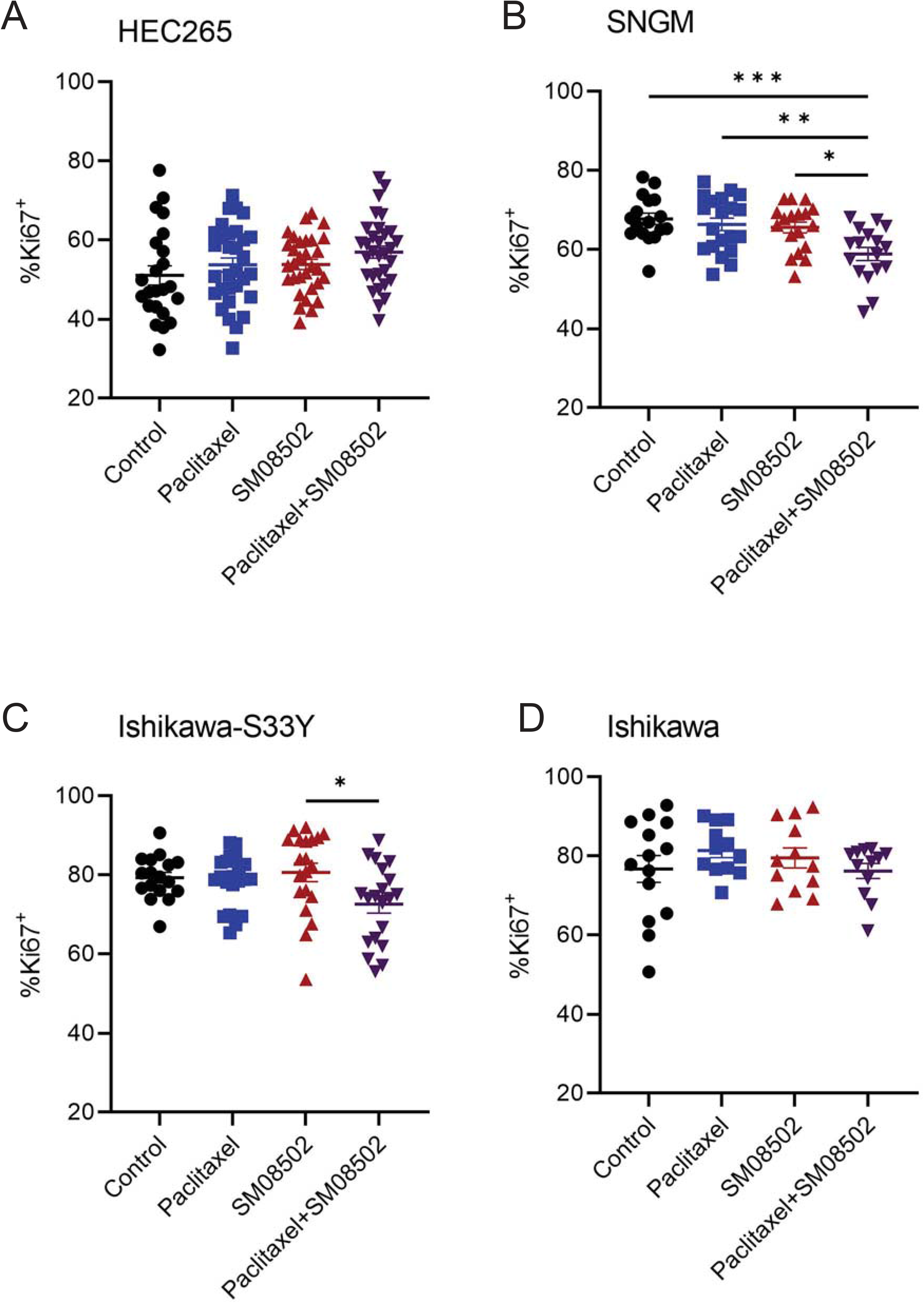
Analysis of tumor proliferation as evaluated by %Ki67+ cells for in-vivo tumor samples taken at time of necropsy. Cell lines evaluated were **(A)** HEC265, **(B)** SNGM, **(C)** Ishikawa-S33Y, and **(D)** Ishikawa.

Toxicity of both the combination and single agent therapies demonstrated excellent tolerability with no variation in body weight (Supplemental Figure 1A), hematologic, or liver function testing at time of necropsy (Supplemental Figure 1B).

### SM08502 increases alternative splicing

In vitro analysis of SM08502 treatment, as shown above, demonstrates decreased Wnt pathway (LCF) transcriptional activity. To evaluate the mechanism of this we evaluated the RNA of control EC cells vs those treated with SM08502 alone. In four independent cell models, a significant increase in alternative splicing (AS) events occurred after treatment (Figure 6).

**Figure 6:**
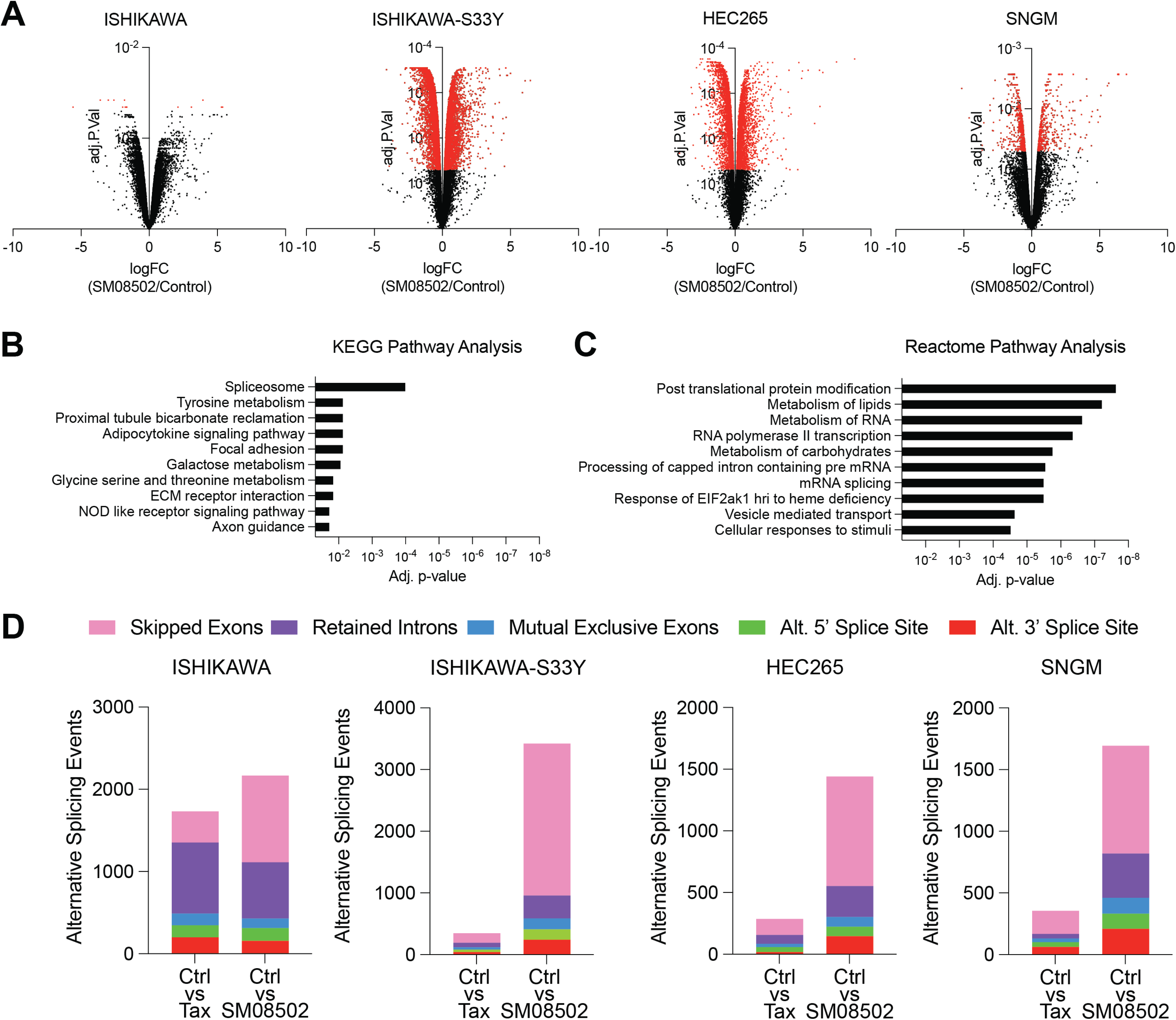
SM08502 significantly reprograms the transcriptome and splicing of endometrial cancer. Four endometrial cancer cell lines (ISHIKAWA, ISHIKAWA-S33Y, HEC265, and HEC108) were treated with vehicle control (DMSO, Control), SM08502 (100 nM for 24 hours), or Taxol (100 nM for 24 hrs). RNA was collected and used for RNA-sequencing (Ribodepleted library preparation with 20 million 150 bp read sequencing). **A)** Volcano plot of differentially expressed genes (red spots, adjusted p<0.05) between control and SM08502. **B)** KEGG Pathway analysis to examine gene enrichments of differentially regulated genes. **C)** Reactome Pathway Analysis to examine gene enrichments of differentially regulated genes. **D)** Replicate Multivariate Analysis of Transcript Splicing analysis of splicing events comparisons in Control versus Taxol and Control versus SM08502. Colors indicate specific splicing events that are significantly (p. adj <0.05) detected compared to control.

Transcriptome analysis via RNA sequencing demonstrates a statistically significant variation in multiple splicing pathways in both the KEGG and Reactome pathway analysis (Figure 6B and 6C). Consistently, measurements of AS events among transcriptome data revealed a significant increase in AS events with the use of SM08502 compared to paclitaxel alone, most notably in skipped exons and retained introns as demonstrated in Figure 6D.

## DISCUSSION

SM08502 is a novel orally available small molecule inhibitor of Cdc2-like kinases (CLK1-4) and the dual specificity tyrosine phosphorylation regulated kinases (DYRK1-4). It is a first in class panCLK/panDYRK inhibitor. CLKs are a member of the CMGC kinase family and have typical kinase structure^50, 51^. CLKs function as both tyrosine and serine /threonine kinases, specifically they autophosphorylate on tyrosine and phosphorylate their protein targets on serine^52, 53^. Hence, CLKs are dual-specific kinases. The CLK family has been linked to a number of cancers including prostate, ovarian, pancreatic, breast, and glioblastoma amongst others^54^. Additionally, the role of CLK family kinases in phosphorylating serine rich splicing factors (SRSFs) is augmented by the DYRK-family kinase activity through phosphorylation of both shared and distinct components of the spliceosome machinery^55^. Thus, pan-inhibition of CLK-and DYRK-family kinases is anticipated to provide maximal suppression of the signal-dependent alternative splice junction selection supporting tumor initiation, progression, and emergence of therapy resistance^56^. To our knowledge, our group was the first to evaluate this mechanism of inhibition in ECs.

Previous evaluation of SM08502 in gastrointestinal cancer models demonstrated a reduction in Wnt pathway gene expression^32^. Our initial publication on stage I, grade 1 endometrioid adenocarcinomas demonstrated that *CTNNB1* is an isolated biomarker for recurrence in a population initially thought to be of low risk for recurrence^28^. We subsequently analyzed the genomic variation of these recurrent tumors and identified the Wnt pathway as significantly variant (Figure 1 A-D), highlighting a potential therapeutic approach. Our previous publication demonstrated that effective inhibition of the Wnt/β-catenin pathway in *CTNNB1*-mutated endometrial tumors occurs with downstream inhibition by decreasing β-catenin/TCF transcriptional activity and cellular proliferation^30^. Reproduction of our previous evaluation of SM04690 with the novel bioavailable form of SM08502 demonstrates remarkable similarity in function. Preclinically, SM08502 demonstrates single agent anti-tumor efficacy in EC cell lines. Cellular death occurs in the nanomolar range across four cell lines tested. We further demonstrate that apoptosis is statistically increased after treatment in three of four cell lines and cellular proliferation is statistically decreased in three of four cell lines as well. Single agent therapy uniformly decreased TCF/transcriptional activity across all cell lines regardless of mutational status. Interestingly, the only cell line in which apoptosis was not increased was the S33Y transduced cell line, Ishikawa-S33Y. Yet, the other *CTNNB1*-mutated cell lines, HEC25 and SNGM, each did demonstrate increased apoptosis with treatment, as did the *CTNNB1*-wildtype cell line, Ishikawa. This infers apoptosis may not be directly related to *CTNNB1* mutation. Mutational status appears to be more correlated with decreased proliferation, consistent with our previously published work^30^, as the only cell line that did not demonstrate decreased proliferation after treatment of SM08502 was the *CTNNB1*-wildtype Ishikawa cell line. Interestingly, this same direct relation did not hold up in Ki67 analysis of the in-vivo tumor models. Proliferation, as evaluated by %Ki67+ cells, was only reduced in the SNGM and Ishikawa-S33Y cell lines with combination therapy compared to single agent SM08502 therapy.

Alternative splicing (AS) is a post-transcriptional mechanism that regulates the translation of mRNA. AS occurs via a variation of splicing methods in which pre-mRNA produce variations of mRNA which therefore translate into diverse proteins and functions. AS is regulated by the spliceosome composed of snRNA and splicing factors^57^. There are currently seven AS events described; skipped exons, retained introns, alternate acceptor site, alternate donor site, alternate terminator, alternate promoter, and mutually exclusive exons^58^. Splicing is a normal physiologic function that allows cells to change their protein production in response to their environment and needs. However, abnormal AS can affect cellular functions such as apoptosis, regulation, and angiogenesis and lead to malignancy. AS has been implicated in tumor proliferation, apoptosis, and metastasis in several tumor types^57, 59, 60^.

AS has demonstrated prognostic effects in EC. Several AS prognostic models have been developed as markers for recurrence and surviva^61–63^. Furthermore, specific splicing factors have been identified to demonstrate worse prognosis. Dou et al. identified that AS of *VEGFA* is directly regulated by the RBM10 in ECs^64^. Popli et al. identified that the splicing factor SF3B1 has an oncogenic role in EC tumorigenesis by regulating KSR2 RNA maturation^65^. It is clear that alteration of AS events has a negative impact on ECs, but more evaluation needs to be conducted.

A whole genome analysis of AS events in EC identified 3 major hub genes in a network of prognostic -related alternative splicing events; RNPS2, NEK2, and *CTNNB1*^63^. SM08502 has previously been identified to decrease mRNA expression of the Wnt genes Axin2, LEF1, MYC, TCF7 and TCF7L2^32^. Our study demonstrates that AS events are significantly increased with treatment of SM08502 compared to treatment with paclitaxel. Specifically, skipped exons and retained introns appear to be the most effected. KEGG and Reactome pathway analysis further demonstrates that significant variation occurs in multiple splicing pathways. Other evaluations have demonstrated the CLK inhibition leads to anti-proliferation, migration, and invasion in other cancers eluding to a similar mechanism as we demonstrate in endometrial cancer^54^. CLK and DYRK inhibition modulate AS by directly phosphorylating splicing factors and increased AS events can lead to the reduced expression of proteins in critical for tumor growth, survival, and resistance.

Our evaluation demonstrates a synergy between the SM08502 and paclitaxel and a rational for further exploration in the clinical setting. Phase I data of this drug as a single agent for advanced solid tumors has shown promising tolerability and efficacy in preliminary results reported on clinical trial NCT03355066^66^. Toxicity assessment of this combination demonstrates no obvious cross toxicities or increased toxicities. Overall, this data provides a strong rationale for combining SM08502 and paclitaxel in EC.

## ACKNOWLEDGMENTS

We acknowledge philanthropic contributions from LeBert-Suess Family Endowed Professorship in Ovarian Cancer Research, Kay L. Dunton Endowed Memorial Professorship in Ovarian Cancer Research, the McClintock-Addlesperger Family, Karen M. Jennison, Don and Arlene Mohler Johnson Family, Michael Intagliata, Mary Normandin, and Donald Engelstad. This work was supported by The Department of Defense (Bitler, OC170228, OC200302, OC200225), The American Cancer Society (Bitler, RSG-19-129-01-DDC), NIH/NCI (Bitler, R37CA261987), The American Association of Obstetricians and Gynecologists Foundation, and The American Board of Obstetrics and Gynecology. The imaging experiments were performed in the Advanced Light Microscopy Core part of Neurotechnology Center at University of Colorado Anschutz Medical Campus supported in part by Rocky Mountain Neurological Disorders Core Grant Number P30 NS048154 and by Diabetes Research Center Grant Number P30 DK116073. The University of Colorado Cancer Center Genomics Shared Resource (RRID:SCR_021984), Flow Cytometry Shared Resource (RRID:SCR_022035), Human Immune Monitoring Shared Resource (RRID:SCR_021985) and Biostatistics and Bioinformatics Shared Resource (RRID:SCR_021983) were partly supported by the NIH (P30CA046934). We further acknowledge Biopslice Therapeutics for supplying SM08502 as well as Carine Bossard, John Hill, Jasgit Sachdev, and Darrin Beaupre for collaboration and assistance with *in vivo* study design. Contents of this paper are the authors’ sole responsibility and do not necessarily represent the official views of our sponsors.

## SUPPLEMENTAL MATERIAL

**Supplemental Table 1:** Cell line characteristics

Abbreviations: ER, estrogen receptor; PR, progesterone receptor.

**Supplemental Figure 1:** Body weight and hematologic/hepatic evaluation. **(A)** Mouse body weights were collected every Monday and Friday during the 21-day study as an indication of drug toxicity. No significant weight loss (i.e. 10% or greater of pre-treatment weight) was observed in any mouse. **(B)** Blood samples were collected from mice via cardiac puncture at the time of death and evaluated for CBC and hepatic panels. Missing data is due to hemolyzed samples. Error bars, SEM.

