## Supplementary figures and images for "Combination CDC-like kinase inhibition (CLK)/Dual-specificity tyrosine-regulated kinase (DYRK) and taxane therapy in *CTNNB1*-mutated endometrial cancer"

### Supplemental Figure 1

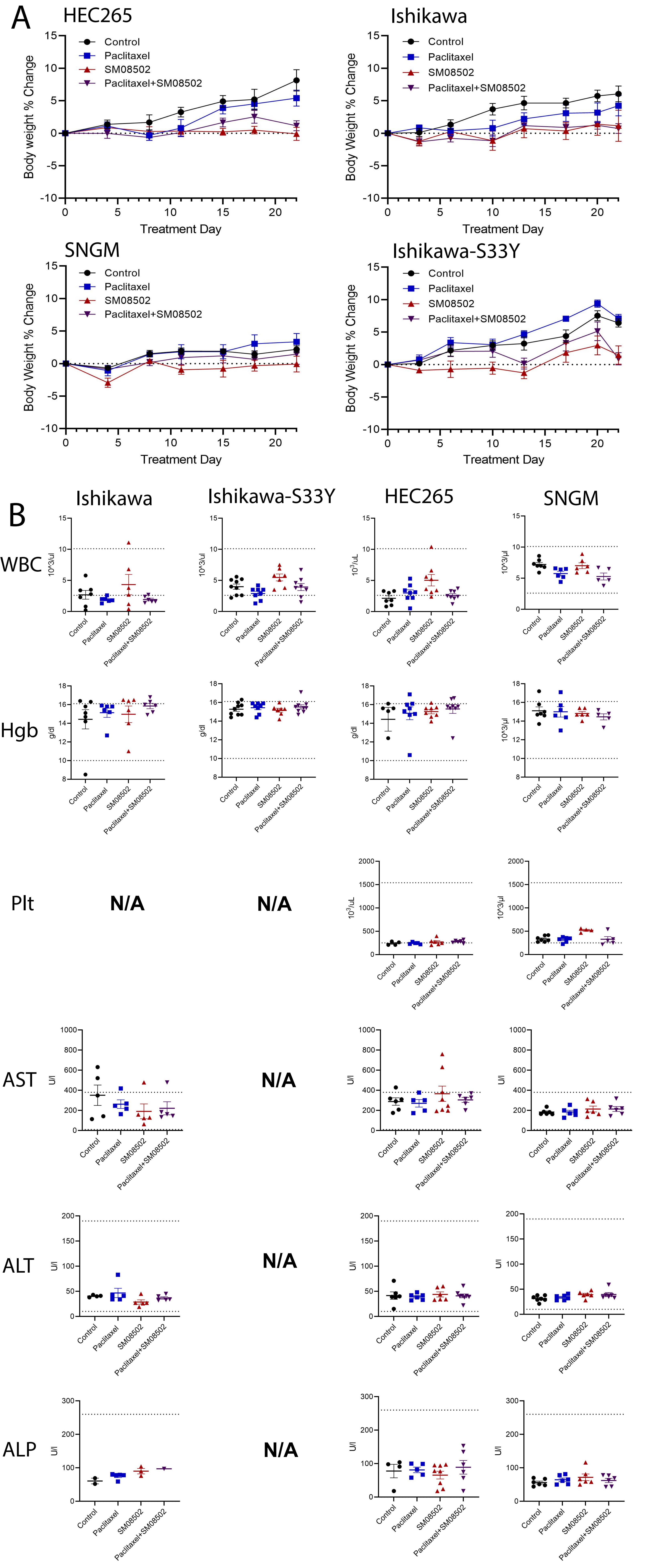
